# Designer grass pea for transgene-free minimal neurotoxin-containing seeds with CRISPR-Cas9

**DOI:** 10.1101/2023.03.26.534271

**Authors:** Tanushree Saha, Ranjana Shee, Salman Sahid, Dibyendu Shee, Chandan Roy, Rajni Sharma, Ashutosh Pandey, Soumitra Paul, Riddhi Datta

**Affiliations:** Department of Botany, University of Calcutta, 35, Ballygunge Circular Road, Kolkata- 700019, West Bengal, India; Department of Botany, Dr. A.P.J. Abdul Kalam Government College, Newtown, Kolkata- 700163, West Bengal, India; Department of Botany, Acharya Prafulla Chandra Roy Government College, Himachal Bihar Phase 1, Matigara, Siliguri, West Bengal, India; National Institute of Plant Genome Research, New Delhi, India

**Keywords:** Grasspea, β-ODAP, *BAHD-AT3*, Cas9-free

## Abstract

Grass pea seeds are consumed as food in several South Asian and Sub-Saharan African nations. However, the presence of the neurotoxic compound N-oxalyl-L-diamino propionic acid (β-ODAP) has restricted its cultivation. Although various cultivars with low β-ODAP levels have been developed, their cultivation is still limited due to the risk of neurolathyrism from long-term grass pea seed ingestion. In this study, we employed the CRISPR-Cas9-mediated gene editing technique to generate grass pea seeds with zero or minimal β-ODAP levels. We targeted the *BAHD-AT3* gene that encodes a key enzyme in the β-ODAP biosynthesis pathway. We developed *bahd-at3* knock-out lines using three gRNAs targeting different regions of this gene and characterized them. Cas9-free independent lines from each event carrying the desired on-target mutation were selected and backcrossed twice with the wild-type to eliminate any off-target mutation present therein. Various agronomical parameters were analyzed from the backcrossed mutant lines and they displayed no phenotypic abnormalities. Interestingly, the seed β-ODAP content ranged between 0.001 % - 0.002 % of dry weight which is 99 % lower than the wild-type. Together, our study reports the development of transgene-free, genome-edited grass peas with insignificant levels of β-ODAP in seeds for safer food in the future.

---

Grass pea (*Lathyrus sativus L*.) is an annual legume crop predominantly cultivated in South Asia and Sub-Saharan Africa (Dixit et al., 2016). Although grass pea seed has high nutritious value, its long-term ingestion causes neurolathyrism in humans and animals because of the presence of an endogenous neurotoxic non-proteinogenic amino acid, β-N-oxalyl-L-α, β-diaminopropionic acid (β-ODAP) (Buta et al., 2019). Several grass pea cultivars with low β-ODAP contents have been developed through conventional breeding, mutagenesis, somaclonal variations, and transgenic approach (Kumar et al.2016, Dixit et al., 2016). Despite these low β-ODAP cultivars, several countries allow limited cultivation of grass peas because their prolonged consumption in large quantities can produce neurolathyrism (Dixit et al., 2016). Therefore, for the safer consumption of grass peas, a cultivar with zero or minimal β-ODAP content (<0.1% seed dry weight) in seeds is required. Although gene editing has been implemented in different legumes, its use in grass peas is limited due to the lack of genetic information (Miller et al., 2022). Therefore, no genome-edited knockout (ko) mutants with reduced β-ODAP content have been reported.

BAHD acyltransferase (BAHD-AT) catalyzes the last step of β-ODAP synthesis by mediating an acyl transfer reaction between oxalyl-CoA and L-α, β-diaminopropionic acid (Goldsmith et al., 2022). Multiple *BAHD-AT* genes have been identified in the grass pea genome. Phylogenetic analysis of these *BAHD-AT* genes revealed that a particular clade has expanded in the legumes significantly. This clade encompasses nine members displaying different tissue-specific expression in grass peas (Edwards et al., 2023). Among them, *BAHD-AT3* displays higher expression over the other isoforms of the clade, with its highest abundance in the late pods (Edwards et al., 2023). Therefore, we hypothesized that loss-of-function mutation of this gene might significantly reduce the β-ODAP content in grass pea seeds. We, therefore, developed *bahd-at3* ko mutants in this study using CRISPR-Cas9-mediated gene editing technology to generate low β-ODAP transgene-free grass pea seeds.

To precisely edit the *BAHD-AT3* (MT457411.1) gene, we designed three different guide RNAs (gRNA1, gRNA2, and gRNA3) by using the CRISPR-P v2.0 (http://crispr.hzau.edu.cn/CRISPR2) and CCTop-CRISPR/Cas9 target online predictor (https://cctop.cos.uni-heidelberg.de:8043/). Three gRNAs (20 bp) targeting different regions of the coding sequence of *BAHD-AT3* with high on-target mutation specificity were selected (Fig. S1). The gRNAs were cloned into the *pHAt-C* vector using Gibson cloning method (Kim et al., 2016). We selected the Indian commercial cultivar, Nirmal, for this study. The recombinant constructs were introduced into this cultivar via *Agrobacterium*-mediated genetic transformation. The putative transformants were screened against hygromycin selection followed by genomic DNA PCR using *hygromycin phosphotransferase* (*hpt)* gene-specific primers (Table S1). Eight independent lines were positive out of fourteen putative transformants generated for gRNA1. For gRNA2, nine independent lines out of fourteen transformants were positive, whereas seven independent lines out of eight transformants were positive for gRNA3 (Fig.S2). Next, we performed PCR-based restriction enzyme (RE) digestion to confirm the on-target mutations in the positive lines of T_1_ generations. *BsaJI* RE was used to digest the PCR products of gRNA1 and gRNA3 ko lines while *BciVI* was used to digest the PCR product of gRNA3 ko lines. Five, eight, and seven independent lines carried on-target mutations for gRNA1, gRNA2, and gRNA3, respectively (Fig.S2). Furthermore, to identify the transgene-free mutant lines, plants were screened using genomic DNA PCR with *Cas9* gene-specific primers (Table S1).

To eliminate the chance of off-target mutations in the ko lines, selected Cas9-free lines from the T_2_ generation were backcrossed twice with WT plants. We further analyzed the on-target mutations in the BC_2_ lines. Eleven BC_2_ lines for gRNA1 and gRNA2 and seventeen BC_2_ lines for gRNA3 carried the on-target mutation (Fig S2). We considered two BC_2_ lines from each set (#1-8-1-2-1-1 and #1-12-2-1-2-5 for gRNA1; #2-8-2-3-2-5 and #2-9-3-1-1-8 for gRNA2; and #3-7-1-5-1-5 and #3-11-1-3-1-5 for gRNA3) for further downstream analysis. Different agronomical traits were analyzed from the selected mutant lines and they displayed no phenotypic alteration compared to the WT plants (Fig 1A-F). Next, to identify the nature of the mutation in the *Cas9*-free mutant lines, we performed PCR-RE based method along with Sanger dideoxy sequencing from the selected lines, and on-target bi-allelic mutations were identified at 3-5 nucleotides upstream of PAM sites (Fig. 1G,H).

**Figure 1.**
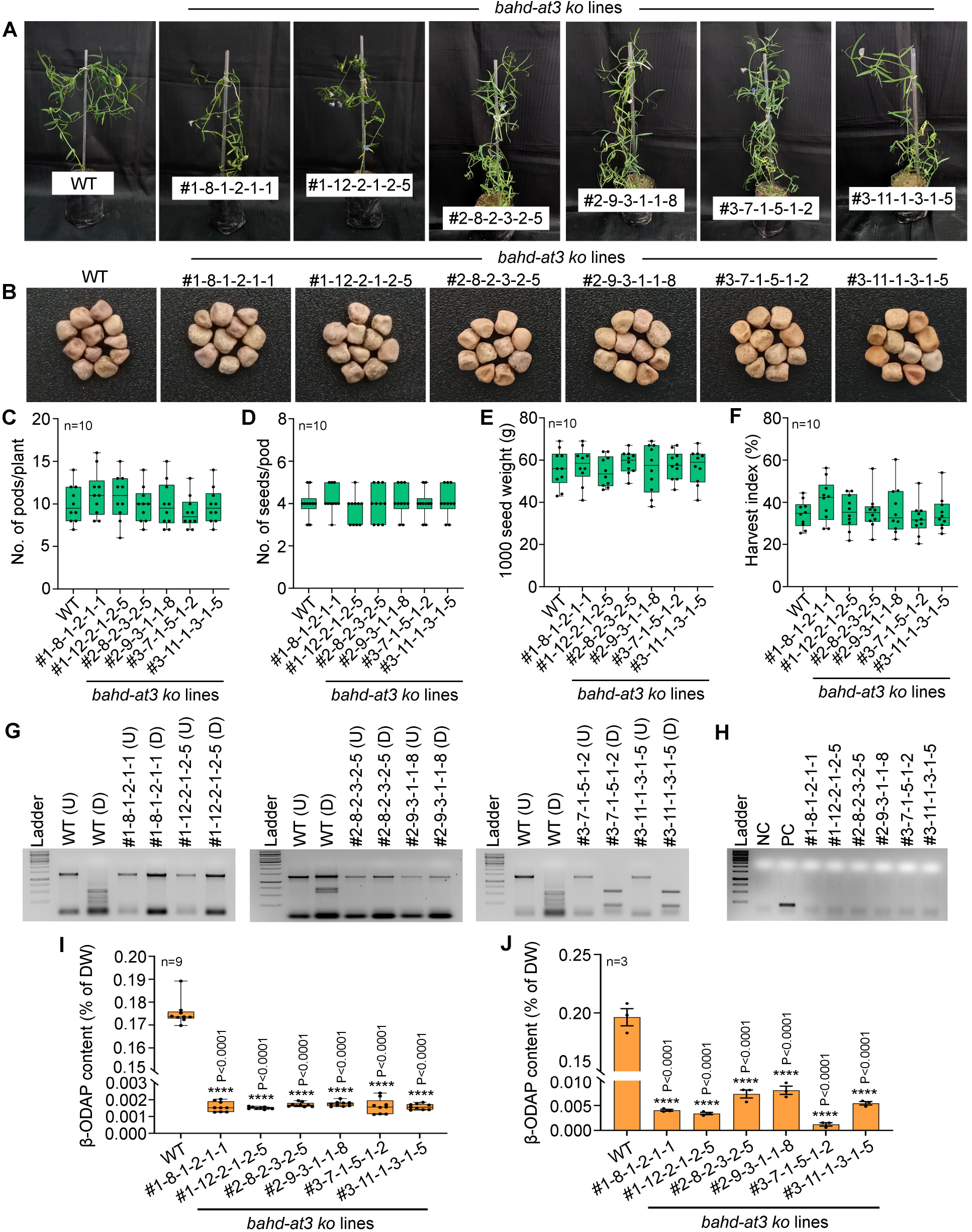
Generation and characterization of bahd-at3 ko lines. Three different gRNAs were designed to target different regions of the *BAHD-AT3* gene and used for CRISPR-Cas9-mediated gene editing of grass pea (Nirmal cultivar). Cas9-free lines T2 carrying the on-target mutation were backcrossed twice to eliminate possible off-target mutation. (A) Representative image of plants from two independent lines generated with each gRNA. (B) Seed morphology, (C) number of pods per plant, (D) number of seeds per pod, (E) 1000 seed weight, and (F) harvest index of the same. (G) Presence of the on-target mutation was confirmed by PCR-RE based method with *BsaJI* for gRNA1 and gRNA3 and *BciVI* for gRNA2. (H) Transgene-free nature of the selected lines was confirmed by PCR with *Cas9*-specific primers. β-ODAP content of the selected lines in (I) seed, and (J) leaf. Statistical differences between the genotypes were analyzed by one-way ANOVA followed by Dunnett’s multiple comparison test and statistical significances were denoted by asterisks and respective P-value in the respective panels. WT: wild-type

Next, we analyzed the β-ODAP content from leaves and mature seeds of WT and ko lines by Ultra Performance Liquid Chromatography (UPLC) method following Ghosh et al. (2015). The β-ODAP content in seeds of the Nirmal cultivar (wild-type) was found to be 0.175 + 0.002 % corroborating the earlier studies that reported 0.08% - 0.33 % of β-ODAP content (Dixit et al., 2016; Edward et al., 2023). Interestingly, the seeds of the transgene-free ko lines displayed a 99 % reduction in β-ODAP content compared with the wild type, while the leaves exhibited around 97 % reduction. The β-ODAP content in the ko lines ranged between 0.001-0.008 % in leaves and 0.002 % in seeds (Fig 1I,J).

In summary, the transgene-free *bahd-at3* ko grass pea lines exhibit up to 99 % lower β-ODAP content while maintaining their original agronomic traits. Together, this study reports the generation of transgene-free gene-edited minimal β-ODAP containing grass peas, which is a big step forward for the crop improvement programs.

## Acknowledgements

This work has been supported by the Science and Engineering Research Board - Women Excellence Award, Government of India [WEA/2020/000009]. We thank Central Instrumentation Facility of Department of Botany, University of Calcutta and Department of Botany, Dr. A. P. J. Abdul Kalam Government College. TS also acknowledge Council of Scientific and Industrial Research (CSIR), Government of India for her fellowship.

## Author Contributions

RD conceived and designed the original research plan; TS and RS performed grasspea transformation, TS performed backcrossing and subsequent phenotyping, SS and DS generated the constructs, CR maintained the plants; RSH and AP performed the metabolic analysis, RD and SP analyzed the data and wrote the manuscript.

## Supporting information

Figure S1. Different gRNAs used in this study

Figure S2. Flowchart for development of *Cas9*-free *bahd-at3* ko lines

Table S1. List of primers used in this study

Appendix S1: Materials and methods

