## Appendix 1 for "Designer grass pea for transgene-free minimal neurotoxin-containing seeds with CRISPR-Cas9"

**Appendix S1: Materials and methods**

*Design of gRNAs, cloning and vector construction*

For CRISPR–Cas9 mediated gene editing in *Lathyrus sativus*, different guide RNAs (gRNAs) specific to *BAHD-AT3* (MT457411.1) were designed using the CRISPR-P v2.0 webserver (http://crispr.hzau.edu.cn/CRISPR2) and CCTop-CRISPR/Cas9 target online predictor (https://cctop.cos.uni-heidelberg.de:8043/). Three gRNAs (20 bp) targeting different regions of coding sequence of *LsBAHD-AT3* with high on-target and low off target mutation specificity were selected (Fig S1). All these gRNAs (gRNA-1, gRNA 2 and gRNA-3)were cloned into the *pHAt-C* vector using Gibson cloning method (Kim et al; 2016). This vector contains Cas9 endonuclease-encoding gene under the control of U6 promoter as well as *hygromycin phosphotransferase* (*hpt*) gene as selection marker. All the constructs were sequenced by dideoxy chain-termination sequencing method (Sanger et al., 1977).

*Generation and screening of bahd at3 knock out lines*

The seeds of grass pea (*L. sativus*) variety Nirmal were procured from Pulses and Oilseeds Research Station, Berhampore, Murshidabad, West Bengal, India. Seeds were surface sterilized by treating with 2% Sodium hypochloride (Merck) and Tween 20 (Merck) followed by repeated washing with sterile distilled water. The surface sterilized seeds were inoculated on Murashige and Skoog (MS) medium for 7 d under 16 h light/ 8 h dark photoperiod for germination (Murashige and Skoog, 1962; Barik et al., 2005). After 7 d, epicotyl segments were infiltrated with *Agrobacterium* strain GV3101 harbouring recombinant constructs, followed by co-cultivation and selection on shoot organogenesis media containing hygromycin for selection as described before (Barik et al., 2005). The regenerated shoots were placed on half-strength MS medium (Hi-Media) for root initiation and transformed plantlets ($T_0$) were established in soil. For each construct, we generated multiple independent lines. The putative knock out (ko) lines of three gRNAs along with wild-type were maintained up to $T_2$ generations in a growth condition at 21±2 ºC, 60% relative humidity, and a photoperiod of 16 h light/8 h dark cycle in plant growth chamber.

*Analysis of cas9 free and on-target mutation of ko lines*

The genomic DNA of *bahd-at3* ko lines from three different events #1,#2, #3 ($T_0$ generation) were extracted by CTAB (Cetyltrimethylammonium bromide) method and genomic DNA PCR was carried out with *hpt* gene-specific primers to screen the transformed plants. The selected transformants were grown for $T_1$ generation. The genomic DNA was isolated from different independent $T_1$ lines and were used for genomic DNA PCR using *Cas9* specific primers to identify *Cas9*-free lines. The selected *Cas9*-free lines were used for on-target (*BAHD-AT3* specific) mutations by PCR-RE based method. The PCR-amplified products of *BAHD-AT3* gene from ko lines along with wild-type were digested with restriction enzyme (RE) *BsaJ*I and *BciVI* (NEB). Furthermore, to identify the specificity of the on-target mutation sequences in ko lines, nested PCR followed by Sanger sequencing was carried out. The DNA sequences were aligned through multiple sequence alignment using CLUSTALW (https://www.genome.jp/tools-bin/clustalw) (Thompson et al., 1994). The Cas9-free ko mutant plants were maintained up to $T_2$ generation.

*Back crossing between ko lines and WT*

To eliminate the chance of off-target mutations in the ko lines, selected Cas9-free ko $T_2$lines were backcrossed twice with wild-type plants. The plants of independent ko lines were emasculated and selected as a female parents while wild-type plants were considered as male parent. The on-target mutations were identified from different back crossing 1 ($BC_1$) lines and selected lines were grown and used for second round of backcrossing. The different $BC_2$ lines were screened for on-target mutations and selected lines were grown for further analysis.

*Evaluation of agronomical traits*

Several agronomical traits namely seed germination percentage, plant height, flower and seed morphology, number of pods per plant, number of seeds per pod, shoot dry weight, harvest index were analysed from $BC_2$ Cas9-free ko lines and compared with the wild-type plants.

*Determination of β-ODAP content*

To analyze β-ODAP content from wild-type and ko lines, Ultra Performance liquid Chromatography was performed using UPLC system with photodiode array detector (Agilent Technologies 1260 Infinity II). Briefly, 100 mg leaves and seeds were harvested, homogenized in 1 ml of 0.1% formic acid and centrifuged (Eppendorf 5810 R, Germany) at 10,000×g for 10 min at 10 °C. After centrifugation, supernatants were passed through 0.22 µm PVDF membrane (Millex®-GV) and 100 µl of each samples were used for quantification. For UPLC analysis, 4 µl of sample was injected with flow rate of 0.05 ml/min. The column temperature was set at 26°C. The sample was applied on reverse phase Zorbax Eclipse plus C18 column (2.1X 55 mm, 1.8 micron). The mobile phase A contained water (Merck) and 0.1% formic acid (Sigma), whereas mobile phase B was acetonitrile (Sigma) containing 0.1% formic acid. The gradient programme was set following Ghosh et al., 2015. The data were calculated by Agilent Open LAB CDS version 2.4.0.695 software (Santa Clara, California, United States). The β-ODAP standard curve was calibrated using β-ODAP standard (Sigma, MO,USA).
