## Supplementary material for "Designer grass pea for transgene-free minimal neurotoxin-containing seeds with CRISPR-Cas9": Figure S1-S2

A

| sgRNA_id | Score | Sequence | Strand | Position | %GC |
|---|---|---|---|---|---|
| sgRNA_1 | 0.7209 | GACCAAGATTGACTCCTCCG**CGG** | + | 885 | 55% |
| sgRNA_2 | 0.7163 | ATCGGAGGTTGGATACTATC**CGG** | - | 594 | 55% |
| sgRNA_3 | 0.5157 | ACTCAATCGATCTAACACCA**TGG** | + | 84 | 40% |

B

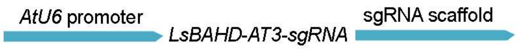


Figure S1. Different gRNAs used in this study

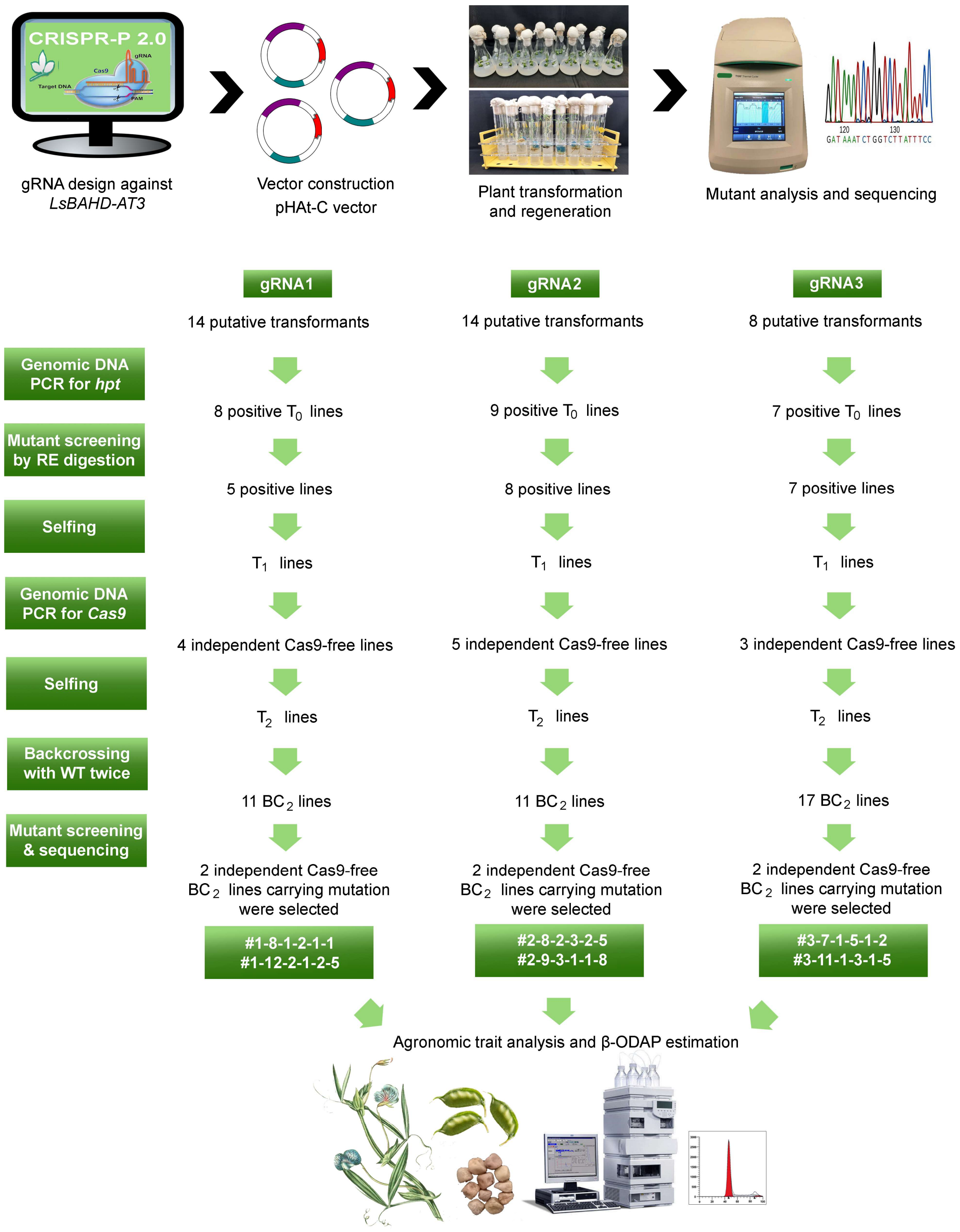


Figure S2. Flowchart for development of *Cas9*-free *bahd-at3* ko lines
