## Supplementary material for "Designer grass pea for transgene-free minimal neurotoxin-containing seeds with CRISPR-Cas9": Table S1

Table S1. List of primers

| Name of gene | Accession number | sequence |
|---|---|---|
| *BAHD-AT3* | MT457411 | Forward: 5'AGTTCCATCCAAATCCTCTCC 3'<br>Reverse: 5'CCAGAAGCAGCATCCATAAAC 3' |
| *Cas9* | KU213971 | Forward: 5'ATCCACCTGTTCACCCTGAC 3'<br>Reverse: 5' GCACGTCGTAGGGGTATACT 3' |
| Nested primer *BAHD-AT3* (gRNA1 region) | MT457411 | Forward: 5' TTGCAAGCGGTTTTCACTCA 3'<br>Reverse: 5' CAAGCCCGCCATCTTCTAAC 3' |
| Nested primer *BAHD-AT3* (gRNA2 region) | MT457411 | Forward: 5'GGAATTTTCATTGGCTGCGC 3'<br>Reverse: 5' CAAACATCGGCTTGCAGAGT 3' |
| Nested primer *BAHD-AT3* (gRNA3 region) | MT457411 | Forward: 5' TCCATCAAAATCCTCTCCACAAC 3'<br>Reverse: 5' TTGAGACGACCTGTGAAGGG 3' |
| *hpt* | AF354046.1 | Forward: 5'GTGCTTGACATTGGGGAGTT3'<br>Reverse:5'GATGTTGGCGACCTCGTATT3' |
